# HaploBlocker: Creation of subgroup specific haplotype blocks and libraries

**DOI:** 10.1101/339788

**Authors:** Torsten Pook, Martin Schlather, Gustavo de los Campos, Manfred Mayer, Chris Carolin Schoen, Henner Simianer

**Author notes:** T. Pook; University of Goettingen, Department of Animal Sciences, Center for Integrated Breeding Research, Animal Breeding and Genetics Group, Albrecht-Thaer-Weg 3, 37075 Goettingen, Germany.

## Abstract

The concept of haplotype blocks has been shown to be useful in genetics. Fields of application range from the detection of regions under positive selection to statistical methods that make use of dimension reduction. We propose a novel approach (“HaploBlocker”) for defining and inferring haplotype blocks that focuses on linkage instead of the commonly used population-wide measures of linkage disequilibrium. We define a haplotype block as a sequence of genetic markers that has a predefined minimum frequency in the population and only haplotypes with a similar sequence of markers are considered to carry that block, effectively screening a dataset for group-wise identity-by-descent. From these haplotype blocks we construct a haplotype library that represents a large proportion of genetic variability with a limited number of blocks. Our method is implemented in the associated R-package HaploBlocker and provides flexibility to not only optimize the structure of the obtained haplotype library for subsequent analyses, but is also able to handle datasets of different marker density and genetic diversity. By using haplotype blocks instead of SNPs, local epistatic interactions can be naturally modelled and the reduced number of parameters enables a wide variety of new methods for further genomic analyses such as genomic prediction and the detection of selection signatures. We illustrate our methodology with a dataset comprising 501 doubled haploid lines in a European maize landrace genotyped at 501’124 SNPs. With the suggested approach, we identified 2’991 haplotype blocks with an average length of 2’685 SNPs that together represent 94% of the dataset.

## Introduction

Over the years, the concept of haplotype blocks has been shown to be highly useful in the analysis of genomes. Fields of application range from population genetics, e.g. the mapping of positive selection in specific regions of the genome (Sabeti et al. 2002, 2007) to statistical applications that make use of dimension reduction (Pattaro *et al.* 2008) and tackle the *p* ≫ *n*–problem (Fan et al. 2014). Existing methods commonly define a haplotype block as a set of adjacent loci, using either a fixed number of markers or a fixed number of different sequences of alleles per block (Meuwissen *et al.* 2014). Alternatively, population-wide linkage disequilibrium (LD) measures (Gabriel et al. 2002; Daly *et al.* 2001; Taliun *et al.* 2014; Kim *et al.* 2017) can be used in the identification process to provide more flexibility of the block size based on local genetic diversity. The methods and software (e.g., HaploView, (Barrett et *al.* 2005)) available for inferring haplotype blocks have become increasingly sophisticated and efficient. Although those approaches to infer haplotype blocks have been proven to be useful, existing methods share some key limitations (Slatkin 2008). In particular, the use of population-wide measures of LD limits the ability of existing methods to capture cases of high linkage characterized by the presence of long shared segments caused by absence of crossing over (typically within families or close ancestry). To illustrate this, consider the following toy example of four different haplotypes: 11111111, 10101010, 01010101, and 00000000. If all four haplotypes have the same frequency in the population, pairwise LD (*r*^2^) of adjacent SNPs is zero and LD-based algorithms would not retrieve any structure. However, in this example, knowledge of the first two alleles fully determines the sequence in the segment.

In this work, we use the term “allele” for a genetic variant. This can be a single nucleotide polymorphism (SNP) or other variable sites like short indels. We use the term “haplotype” for the sequence of alleles of a gamete, and not as often done as a short sequence of alleles. Lastly, a combination of multiple adjacent alleles is here referred to as an “allelic sequence”.

As the starting point of our approach (“HaploBlocker”), we assume a set of known haplotypes which can be either statistically determined as accurately phased genotypes, or observed via single gamete genotyping from fully inbred lines or doubled haploids. When the interest is in inferring the longest possible shared segment between haplotypes, a common approach is to identify segments of identity-by-descent (IBD). A tool for the identification of IBD segments is BEAGLE (Browning and Browning 2013), among others. Since IBD is typically calculated between pairs of individuals, a screening step is used to identify haplotypes that are shared by multiple individuals, e.g. by the tool IBD-Groupon (He 2013). A method to detect IBD segments directly for groups of individuals has been proposed by Moltke *et al.* (2011), but is not applicable to datasets with hundreds of haplotypes due to limitations of computing times. A further difficulty is that common methods are not robust against minor variation, leading to actual IBD segments being broken up by calling errors (0.2% with the later used Affymetrix Axiom Maize Genotyping Array (Unterseer *et al.* 2014)) and other sources of defects.

The imputation algorithm of BEAGLE uses a haplotype library given by a haplotype cluster (Browning and Browning 2007). The haplotype library in BEAGLE, which is used to initialize a Hidden Markov Model for the imputing step, is only given in a probabilistic way. This means that there are no directly underlying haplotype blocks that could be used for later statistical application.

Our goal is to provide a conceptualization of haplotype blocks that can capture both population-wide LD and subgroup-specific linkage, and does not suffer from some of the limitations of IBD-based methods. Unlike common definitions that consider haplotype blocks as sets of adjacent markers, we define a haplotype block as an allelic sequence of arbitrary length.

Haplotypes with a similar sequence are locally assigned to the same block. Haplotype blocks are subgroup specific, so that a recombination hot spot appearing in a subgroup of haplotypes does not affect the boundaries of other blocks. This leads to very long blocks, which can cover the same region of the genome, but may vary in the allelic sequence they represent. Even an overlap between the allelic sequences represented by different haplotype blocks is possible.

Subsequently, we start with a large set of identified haplotype blocks and reduce this set to the most relevant blocks and thus generate a condensed representation of the dataset at hand. We define this representation as a haplotype library and, depending on the topic of interest, selection criteria for the relevance of each block can be varied appropriately to identify predominantly longer blocks or focus on segments shared between different subpopulations. So, the standard input of HaploBlocker is a phased SNP-dataset and the output is a haplotype library that in turn can be used to generate a block dataset. A block dataset contains dummy variables representing the presence/absence of a given block (0 or 1) or, in case of heterozygotes, a quantification of the number of times (0, 1 or 2) a block is present in an individual. The usage of haplotype blocks instead of SNPs allows the use of a variety of new methods for further genomic analyses since the number of parameters is usually massively reduced and haplotype blocks provide a natural model for local epistatic interactions.

## Materials and Methods

The aim of HaploBlocker is to represent genetic variation in a set of haplotypes with a limited number of haplotype blocks as comprehensively as possible. The main idea of our method is to first consider short windows of a given length and increase the length of the analyzed segments in an iterative procedure involving the following steps:

- Cluster-building
- Cluster-merging
- Block-identification
- Block-filtering
- Block-extension
- Target-coverage (optional)
- Extended-block-identification (optional)

Before we elaborate on each step in the following subsections, we give an outline of the three major steps. For a schematic overview of HaploBlocker, we refer to Figure 1. In the first step, we derive a graphical representation of the dataset (“window cluster”) in which a node represents an allelic sequence and an edge indicates which and how many haplotypes transition from node to node (cluster-building). As locally similar allelic sequences are grouped together, this step also handles robustness against minor deviations (e.g. calling errors). In the second major step, we identify candidates for the haplotype library based on the window cluster. We call this step block-identification and use it to generate a large set of haplotype blocks. In the third and last major step (block-filtering), the set of candidates is reduced to the most relevant haplotype blocks and thereby the haplotype library is generated. In addition to specifying the physical position of each block, we have to derive which haplotypes are included. The fact that blocks are subgroup specific makes the identification of the most relevant blocks complicated so that we split this task into two separate, but closely connected steps (block-identification and block-filtering).

**Figure 1.**
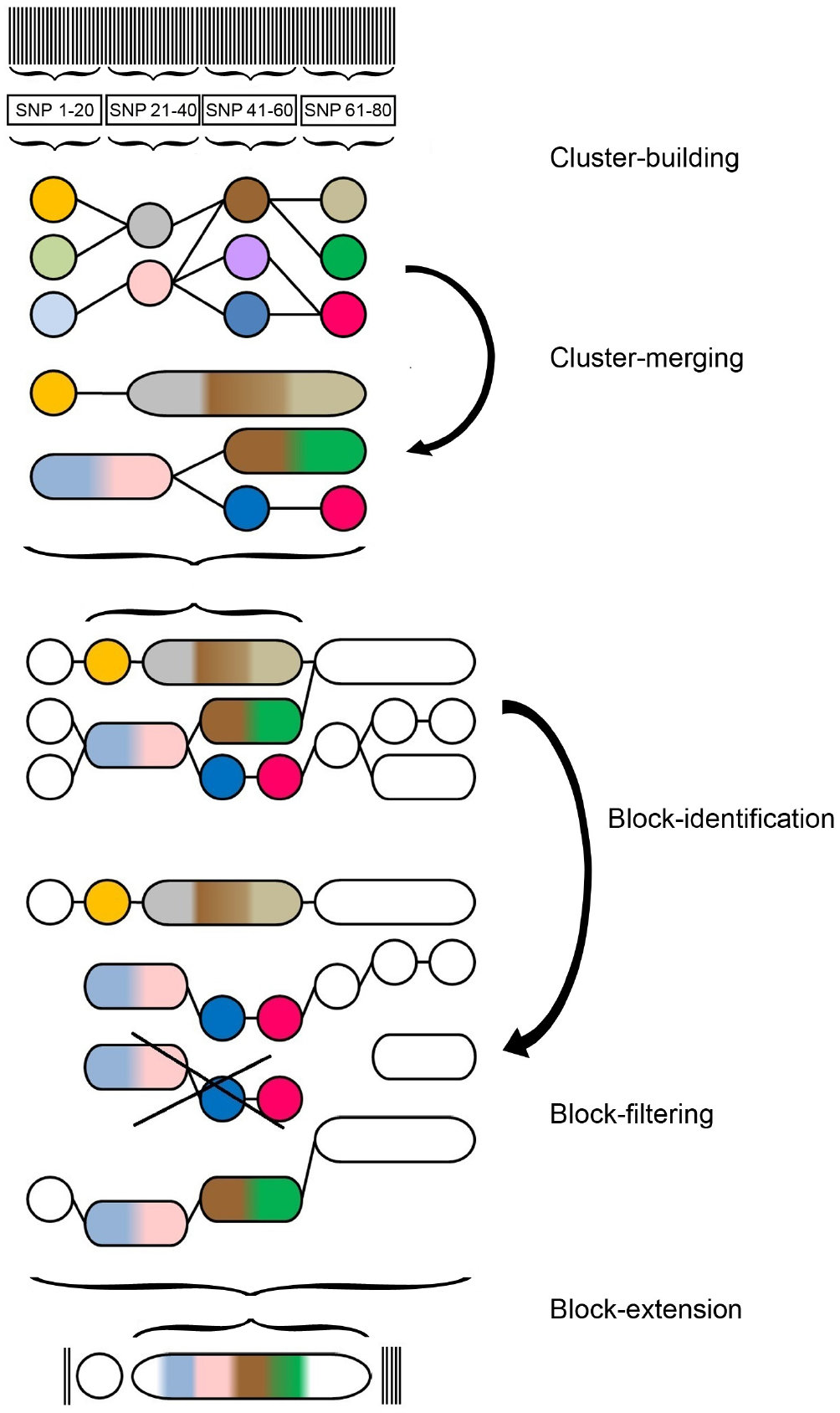
Schematic overview of the steps of the HaploBlocker method: (1) Cluster-building: Classifying local allelic sequences in short windows into groups. (2) Cluster-merging: Simplifying window cluster by merging and neglecting nodes. (3) Block-identification: Identifying blocks based on transition probabilities between nodes. (4) Block-filtering: Creating a haplotype library by reducing the set of blocks to the most relevant ones for the later application. (5) Block-extension: Extending blocks by single windows and SNPs. The same allelic sequences in different steps are coded with the same colors in the graph.

Minor steps in our procedure are cluster-merging and block-extension. The former reduces the computing time in the subsequent steps, whereas the latter increases the precision of the result. However, neither step has a major impact on the resulting haplotype library. Since various parameters are involved in the procedure, their value might be chosen by means of an optimization approach and/or a dataset can be processed with multiple parametrizations in the cluster-building, cluster-merging and block-identification-step. For more details, we refer to the subsections on target-coverage (Supplementary Material File S1) and extended-block-identification.

The next subsections deal with the graphical depiction of the haplotype library and the information loss incurring through the suggested condensation of genomic data. Subsequently, we discuss possible applications, namely the power of our method to recover founder haplotypes of a population and a block-based version of extended haplotype homozygosity (EHH, (Sabeti *et al.* 2002)) & integrated extended haplotype homozygosity (IHH, (Voight *et al.* 2006)). In the last subsection, we introduce the datasets used in this study. Our method, as well as all discussed applications, are available for users by the correspondent R-package HaploBlocker (R Core Team 2017; Pook and Schlather 2018). The default settings of the arguments in the R-package correspond to the thread of the following subsections.

### Cluster-building

In the first step of HaploBlocker we devide the genome into non-overlapping small windows of size 20 markers as a default value. Accordingly, each haplotype is split into short allelic sequences. To account for minor deviations, we merge groups with similar allelic sequences. For a fixed window, different allelic sequences are considered successively based on their frequency, starting with the most common one. In case a less common allelic sequence differs only in a single marker, they are merged to a group. The allelic sequence of a group (“joint allelic sequence”) in each single marker is the most common allele of all contained haplotypes. Usually this will be the most frequent allelic sequence but when allowing for more than one deviation per window this is not necessarily the case anymore. As a toy example, consider a group containing 4x 11111, 3x 10110, 2x 00111 with a resulting joint allelic sequence of 10111. This robustness against errors may lead to actually different haplotypes to be grouped together. In later steps, we will introduce methods to split these haplotypes into different blocks if necessary. The choice of 20 markers as the window size and a deviation of at most one marker as a default is not crucial and should not have a major effect as long as the window size is much smaller than the later identified haplotype blocks. We will present ways to use flexible window sizes in the extended-block-identification-step. As an example consider a SNP-dataset with 200 haplotypes and 5 markers, given in Table 1. The two most common sequences form separate groups (00011 & 11111). For graphical reasons in later steps, we assign 11111 to group 3 even though it is the second group created. The next allelic sequence (11110) is assigned to the group of 11111, as it is only different in a single allele and the joint allelic sequence remains 11111. In case an allelic sequence could join different groups, it is added to the group containing more haplotypes. Based on the groupings we are able to create a graph, called window cluster (Figure 2, top graph). Here, each node represents a group (and thus a joint allelic sequence) and the edges indicate how many of the haplotypes of each node transition into which adjacent node.

**Table 1.** Exemplary dataset of allelic sequences and their assignment according to the cluster-building-step.

| Frequency | Allelic sequence | Group |
| --- | --- | --- |
| 101 | 00011 | 1 |
| 54 | 11111 | 3 |
| 40 | 11110 | 3 |
| 3 | 10011 | 1 |
| 2 | 01001 | 2 |

**Figure 2.**
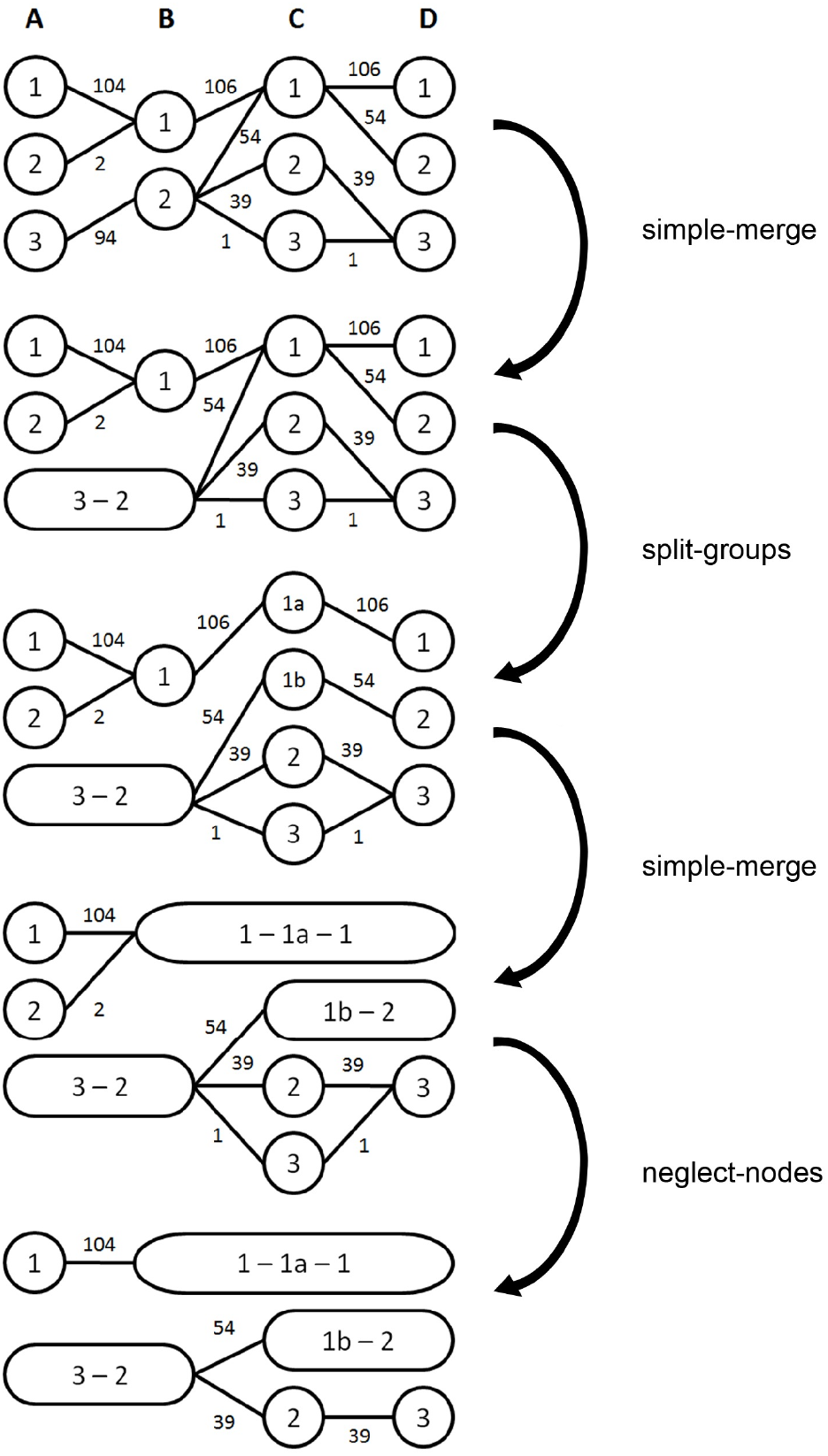
The four parts of the cluster-merging-step. Haplotype frequencies in the window A are according to the toy example given in Table 1.

### Cluster-merging

A window cluster can be simplified without losing any relevant information for later steps of the algorithm. We apply three different techniques:

- simple-merge (SM): Combine two nodes if all haplotypes of the first node transition into the same adjacent node and no other haplotypes are in the destination node.
- split-groups (SG): Split a node into two if haplotypes from different nodes transition into the same node and split into the same groups afterwards.
- neglect-nodes (NN): Remove a node from the cluster if it contains a very small number of haplotypes, 5 say. These removed haplotypes are still considered when calculating transition probabilities between nodes in later steps.

Since the only loss of information in this step stems from neglecting nodes, we first alternately apply SM and SG until no further changes occur. Next, we apply the sequence of NN, SM, SG, SM until fixation of the window cluster. We neglect rare nodes, since a block with few haplotypes (in the most extreme case a block with one haplotype over the whole genome) does not reflect much of the population structure and would have little relevance for further genomic analyses. It should be noted that the minimum number of haplotypes per node in NN does not depend on the number of haplotypes in the sample. This is mainly done to ensure a similar structure of the later obtained haplotype library when adding haplotypes from a different sub-population. Nevertheless the option to modify this parameter is given, in case one is mostly interested in more common or even rarer allelic sequences.

As an example for the cluster-merging-step consider a dataset with four windows and five different sequences of groups (104x 1111, 54x 3212, 39x 3223, 2x 2111, 1x 3233, Figure 2). Haplotypes in the first window are chosen according to Table 1. In the first step nodes A3 and B2 are merged by SM. Next, node C1 is split into two nodes via SG. This triggers additional SMs (B1-C1a-D1 and C1b-D2). Afterwards, no SM or SG are possible anymore and NN is performed removing A2 and C3. No further SM or SG are possible after this. Consider that even though D3 is the only node following C2 no SM is possible because removed haplotypes are still considered in later transition probabilities and therefore D3 contains one more haplotype than C2.

### Block-identification

In the third step of HaploBlocker we identify the haplotype blocks themselves. As a haplotype block in HaploBlocker is defined as a common allelic sequence in an arbitrarily large window, we use common sequences of nodes in the previously obtained window cluster as a first set of haplotype blocks. The identification process itself is performed by using each node as a starting block. The boundaries of each starting block are given by the boundaries of the node and the allelic sequence is derived by its joint allelic sequence. A block is iteratively extended if at least 97.5% of the haplotypes in a block transition into the same node; deviating haplotypes are removed. Haplotypes filtered out in this step can rejoin the block if their allelic sequence matches that of the joint allelic sequence of the final haplotype block in at least 99% of the markers. The joint allelic sequence is derived by computing the most common allele in each marker for the contained haplotypes. The choices of 97.5% and 99% worked well in our tests, but any value close but not equal to 100% should work here. This again allows the user some flexibility in how long (in terms of physical length) the haplotype blocks should be and how different jointly considered haplotypes are allowed to be. In a similar way, each edge of the window cluster is used as a starting block. Here, boundaries are given by the boundaries of the two connected nodes. The haplotype blocks identified here will not all be part of the final haplotype library but instead are just a set of candidates from which the most relevant ones will be selected in the block-filtering-step. Note that the share of allowed deviations in this step (1%) is lower than in the cluster-building (1 of 20 markers − 5%), since the size of the identified segment is longer than a single window and the total number of deviations should get closer to the expectation (Unterseer *et al.* 2014).

To illustrate the method, consider an excerpt of a window cluster given in Figure 3. Nodes 2, 3, 4 represent the sequence of groups 3223 of Figure 2. When considering the second node as a starting block, we cannot extend the block because there are multiple possible nodes for the contained haplotypes (beforehand: nodes with 88 (93.6%) and 6 (6.4%); afterwards: 54 (57.4%), 1 (1.1%), 39 (41.5%)). When using the fourth node of the excerpt, the block can be extended till the second and fifth node of the cluster since 39 of the 40 haplotypes transition (97.5%) into the same adjacent node. One ends up with the same block when using the third node or the edges including 39, 39 and 40 haplotypes. In case all included haplotypes transition into the same node in the first window, the block could be extended even further. Note that in this step different allelic sequences of the same node (cf. cluster-building-step) can be in different haplotype blocks if they transition into different nodes in later steps (e.g 11111 (54) and 11110 (39+1) in the first window (Table 3 & Figure 2). For an extension to further increase the size of the set of haplotype blocks, we refer to the extended-block-identification-step.

**Figure 3.**
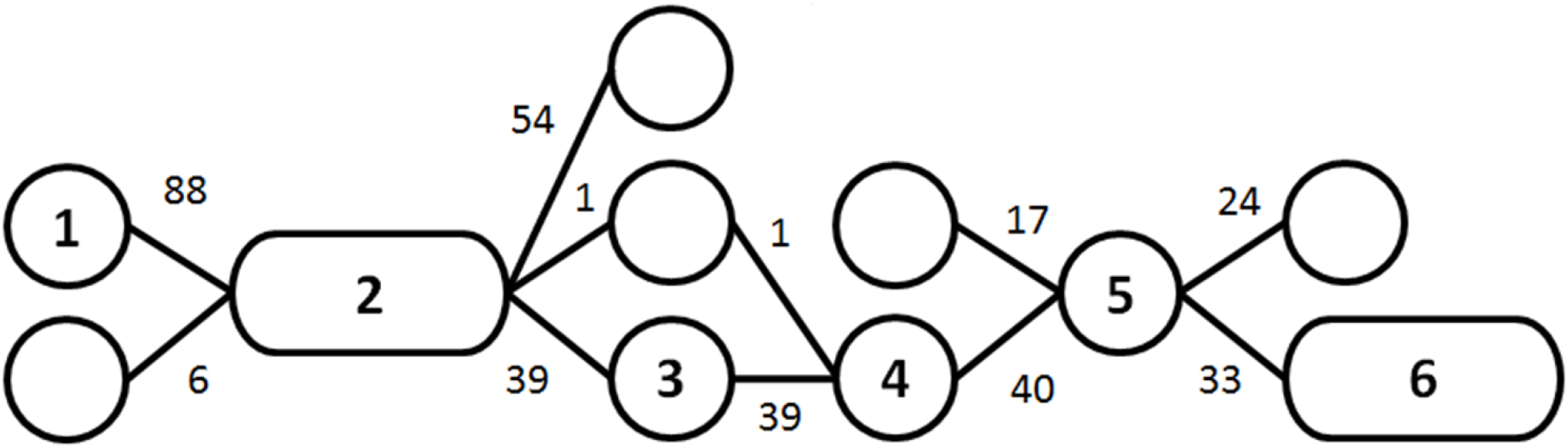
Excerpt of a window cluster. This included all edges (transitions) from the nodes of one of the common paths in the example dataset.

### Block-filtering

After the derivation of candidates, we reduce the set of all haplotype blocks to a haplotype library of the most relevant blocks to represent a high proportion of the dataset with a small number of blocks. First, we compute a rating *r_b_* for each block *b* that depends on its length (*l_b_*) and the number of haplotypes (*n_b_*) in it:

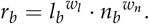

Here *w_l_* and *w_n_* represent weighting factors with default values *w_l_* = 1 and *w_n_* = 1. Note that only the ratio between both parameters matters.

Based on these ratings we determine which haplotype block is the most relevant in each single cell of the SNP-dataset matrix. Iteratively, the blocks with the lowest number of cells as the most relevant block are removed from the set of candidates, until all remaining blocks are the most relevant block in a given number of cells. For this, we will later use the abbreviation MCMB (<u>m</u>inimum number of <u>c</u>ells as the <u>m</u>ost relevant <u>b</u>lock). For our datasets, 5’000 was a suitable value for MCMB but without prior information, we recommend to set a target on what share of the SNP-dataset is represented by at least one block (“coverage”). For details on the fitting procedure we refer to the Supplementary Material (File S1). In case of our example given in Figure 3 we end up with block *b*_1_ (green in Figure 4) including 94 haplotypes ranging from window 2 to 3 (node 2) with a rating *r*_*b*_1__ = 940 and block *b*_2_ (red in Figure 4) ranging from window 2 to 6 (nodes 2,3,4,5) with a rating *r*_*b*_2__ = 975. To simplify the example, we assume that no other blocks have been identified. *b*_2_ has a higher rating, therefore cells containing both blocks are counted as cells with *b*_2_ as the most relevant block. This leads to b1 having 550 cells of the SNP-dataset as the most relevant block and *b*_2_ having 975.

**Figure 4.**
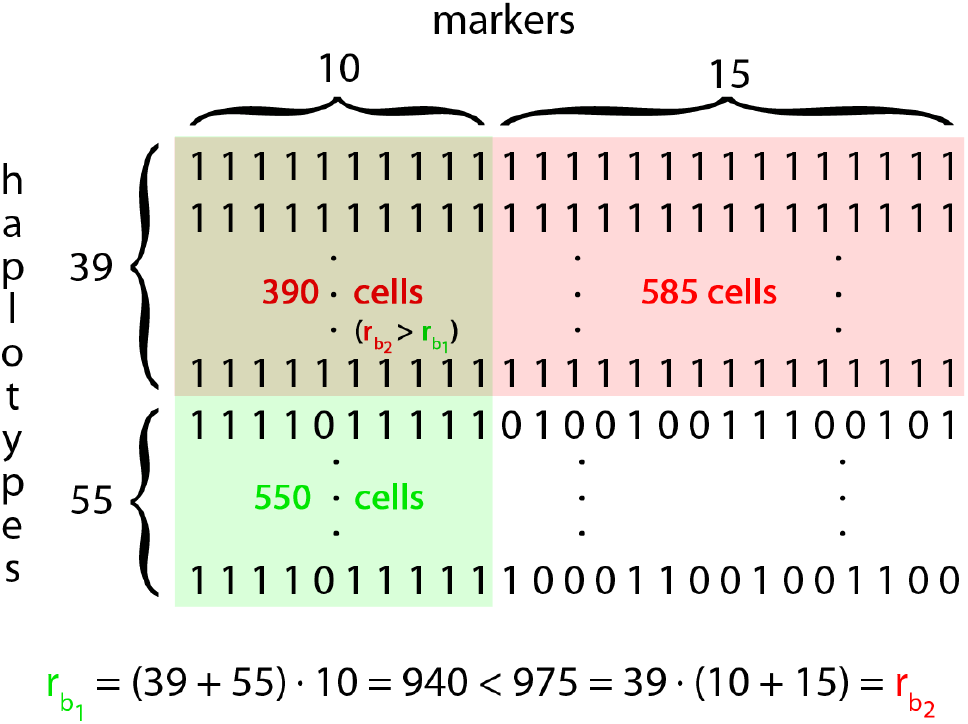
Toy example for the calculation in the block-filtering-step with *w_l_* = *w_n_* = 1.

It has to be noted here that the blocks in the final haplotype library can overlap. In case the MCMB is 550 or smaller, overlap occurs in our example and typically can be observed when a short segment is shared in the majority of the population and a smaller subgroup shares a longer segment which includes the short segment. This will lead to dependencies in the presence/absence of blocks that can be addressed in a similar way as linkage disequilibrium between markers.

Even if *w_l_* or *w_n_* is set to zero, there is still an implicit weighting on both the length and the number of haplotypes since each haplotype block has to cover a certain number of cells of the SNP-dataset (MCMB). The overall effect of *w_l_* and *w_n_* is higher when more candidates were created in the block-identification-step.

### Block-extension

The haplotype blocks that have been identified in the previous step are limited to the boundaries of the nodes of the window cluster. Haplotypes in the blocks will transition into different adjacent nodes since the block was previously not extended (cf. block-identification). Nevertheless, different nodes can still have the same allelic sequence in some adjacent windows.

First, haplotype blocks are extended by full windows if all haplotypes are in the same local group (cf. cluster-building) in the adjacent window. If the haplotypes of a specific block transition into different groups in the adjacent window, the block is still extended if there is no variation in the following 20 windows. By doing this, we account for possible errors that could have been caused by translocations or phasing issues, for instance. The choice of 20 windows is again rather arbitrary and should be chosen according to the minimum length of the blocks one is interested in. In any case, all SNPs with variation in a block are identified and reported in the outcome as a possible important information for later analyses.

Secondly, blocks are extended by single adjacent SNPs following similar rules as the window extension. As a default, we do not allow for any differences here since haplotypes in the block must have some difference in the adjacent window. In case of working with a large number of haplotypes and aiming at identifying the exact end of a block, one might consider allowing for minor differences.

### Extended-block-identification (optional)

When extending a haplotype block in the block-identification-step, haplotypes transitioning into a different node are removed. Instead, one could consider both the short segment with all haplotypes and the long segment with fewer haplotypes. As the number of candidates is massively increased, it is recommended to consider the long segment only when at least a share *t* of haplotypes transition into that node. In our tests *t* = 0.95 was a suitable value for this. Overall, this procedure will lead to the identification of even longer haplotype blocks as candidates for the haplotype library.

To obtain even more candidates in the block-identification-step, one might compute multiple window clusters under different parameter settings (especially concerning window sizes). This provides additional robustness of the method. Especially in case finally obtained haplotype blocks are short, the relevant haplotype blocks can only be identified with a low window size in the cluster-building-step.

Both extensions require substantially more computing time and thus are not included in the default settings of the associated R-package HaploBlocker (R Core Team 2017; Pook and Schlather 2018). The R-package contains an adaptive mode using window sizes of 5,10,20,50 markers and a target coverage of 90%.

### Graphical representation of haplotype blocks

We suggest a graphical representation of a haplotype library to display transition rates between blocks in analogy to bifurcation plots (Sabeti *et al.* 2002; Gautier and Vitalis 2012). This requires ordering of the haplotypes according to their similarity in and around a given marker. For technical details on the sorting procedure we refer to the Supplementary Material (File S2).

### Assessment of information content of haplotype blocks

HaploBlocker provides a condensed representation of the genomic data. We next discuss how to quantify the amount of information lost in the process of condensing a SNP-dataset to a block dataset. At a recent conference, de los Campos (2017) proposed three methods for estimating the proportion of variance of an omics set (e.g. high-dimensional gene expression data, methylation or markers) that can be explained by regression on another type of omics data. We used a modified version of the second method proposed by de los Campos (2017) to estimate the proportion of variance of the full SNP-set genotypes that can be explained by a regression on the blocks of a haplotype library. For the computations in this work the R-packages sommer (R Core Team 2017; Covarrubias-Pazaran 2016) and minqa (Powell 2009) were used with overall very similar results. The methodology can be briefly described as follows:

In traditional SNP-based genomic models (Meuwissen *et al.* 2001), a phenotype (*y*) is regressed on a SNP-dataset (*X*) using a linear model. Entries in *X* are usually 0,1,2 with dimensionality being the number of individuals (*n*) times the number of markers (*p*).

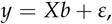

assuming that the markers only have additive effects *b*. Hence, the vector of genomic values *g* = *Xb* is a linear combination of the SNP genotypes. In order to estimate the proportion of *g* explained by the haplotype library, we regress the genomic values *g* onto the block dataset represented by a *n × q* matrix Z, say, of entries 0,1,2. Here *q* is the number of blocks, with *q* usually being much smaller than *p*:

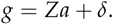

From this perspective, genomic prediction based on haplotype blocks searches for a vector *Za* that is optimal in some sense. For instance, in ridge regression, such a vector is obtained by minimizing a penalized residual sum of squares. It has to be noted that *ε* is an error term that includes non-genetic effects whereas *δ* is an error term resulting from genetic effects that cannot be explained by additive effects (*a*) of single blocks. In random effect models the proportion of the variance of *g* explained by linear regression on the haplotype library can be estimated using either Bayesian or likelihood methods like REML (Patterson and Thompson 1971). This proportion of variance explained will vary from trait to trait. We estimate the distribution of the proportion of variance of “genomic vectors” (i.e., linear combinations of SNP genotypes) using a Monte Carlo method. The method proceeds as follows:

1. Sample a vector of weights (*b_s_*) completely at random (e.g. from a standard Gaussian distribution)
2. Compute the underlying genomic value by forming the linear combination: *g_s_* = *Xb_s_*
3. Estimate the proportion of variance of *g_s_* that can be explained by regression on haplotype blocks
4. Repeat 1.- 3. for a large number of random vectors *b_s_*

In contrast to commonly used methods like canonical correlation (Witten et al. 2009), this method is asymmetric in that it leads to different results by switching the roles of *X* and *Z*. The underlying genomic value is then generated based on the block dataset (*g_s_* = *Zb_s_*) and regressed on the SNP-dataset *X*. Since we compute the share of the variance of one dataset explained by the other dataset, the share of variation that is not explained can be interpreted as previously underused information. An example for underused information are local epistatic interactions that can be modeled via a block but are usually not fully captured by linear regression on single markers.

Recent work has suggested that the direct estimation of the heritability using REML variance components is biased (Schreck and Schlather 2018), so we use their proposed estimator. For the traditional estimates using REML estimates as proposed in the conference talk by de los Campos (2017) we refer to the Supplemental Material (Table S1). Overall, results were similar.

### Recovering founder haplotypes

HaploBlocker does not require or make use of pedigree or founder haplotypes, but rather provides a method to recover haplotypes from the founders (or a genetic bottleneck) just based on a genetic dataset of the current generation. To evaluate the ability to recover founder haplotypes, we simulated the generation of a multiparent advanced generation intercross population (MAGIC) based on the breeding scheme given in Zheng et al. (2015). Simulations were performed with the R-package MoBPS (R Core Team 2017; Pook 2018) (available at https://github.com/tpook92/MoBPS) with 19 founding haplotypes, intercrossing with a diallel design, four generations of random mating and ten generations of self-fertilization (Zheng et al. 2015). Each generation contains 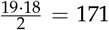 offspring. Genotypes of founders were assumed to be fully homozygous with uniformly distributed minor allele frequencies and under the assumption of equidistant markers (50k markers, 1 chromosome with a length of 3 Morgan, mutation rate of 10^-4^ in each marker). The haplotype phase of the final generation of offspring was assumed to be known.

### Block-based EHH & IHH

The extended haplotype homozygosity statistic (EHH, (Sabeti et al. 2002, 2007)) is defined as the probability of a segment between two markers to be in IBD and can be estimated as:

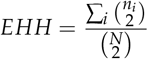

Here *N* is the total number of haplotypes and *n_i_* is the number of occurrences of a given allelic sequence between the markers. In a second step, IHH (Voight *et al.* 2006) for a single marker is defined as the integral when calculating EHH of that marker to adjacent markers (until EHH reaches 0.05).

This concept can be extended to an EHH that is based on haplotype blocks (bEHH). Instead of calculating EHH for each marker, segments between the block boundaries (*a*_1_, *a*_2_, *a*_3_,…) of haplotype blocks are considered jointly. Here *a_i_* is a physical positions (e.g. in base pairs) in the genome. The set of block boundaries contains all start points of blocks, as well as, all markers directly after a block (and not the end point itself). bEHH between segments [*a_i_*, *a*_*i*+1_ − 1] and [*a_j_*, *a*_*j*+1_ − 1] is then defined as the probability of two randomly sampled haplotypes to belong to the same haplotype block, or at least a block with the same allelic sequence in the window [*a_i_*, *a*_*j*+1_ − 1] (with *i* ≤ *j*). bEHH between two markers is set equal to bEHH between the two respective segments. IHH and derived test statistics like XP-EHH or iHs (Sabeti *et al.* 2007) can then be defined along the same lines as with single marker EHH. For a toy example on the computations necessary to compute EHH and bEHH we refer to the Supplementary Material (File S3 & Figure S1).

Overall, bEHH can be seen as an approximation of EHH. Computing times are massively reduced since bEHH scores only need to be computed between pairs of segments instead of SNPs, overall leading to 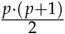 necessary computations, with *p* being the number of segments and SNPs, respectively. Secondly, only allelic sequences of different haplotype blocks, instead of individual haplotypes between the two segments need to be compared for each calculation of bEHH.

As minor deviations from the joint allelic sequence of a haplotype block are possible, the usage of bEHH also provides robustness against calling errors and minor deviations.

### Genotype data used

We applied HaploBlocker to multiple datasets from different crop, livestock and human populations. In the following, we report results obtained with a dataset of doubled haploid (DH) lines of two European maize (*Zea mays*) landraces (*n* = 501 Kemater Landmais Gelb (KE) & *n* = 409 Petkuser Ferdinand Rot (PE)) genotyped with the 600k Affymetrix^®^ Axiom Maize Geno-typing Array (Unterseer *et al.* 2014) containing 616’201 markers (609’442 SNPs and 6’759 short indels). Markers were filtered for being assigned to the best quality class (PolyHighResolution, (Pirani *et al.* 2013)) and having a callrate of 90% or higher. Since we do not expect heterozygous genotypes for DH lines, markers showing an excess of heterozygosity might result from unspecific binding at multiple sites of the genome. Thus, markers were also filtered for having less than 5% heterozygous calls. This resulted in a dataset of 501’124 usable markers. The remaining heterozygous calls of the dataset were set to NA and imputed using BEAGLE 4.0 (Browning and Browning 2016) with modified imputing parameters (buildwindow=50, nsamples=50, phase-its=15).

Secondly, we used a dataset containing *n* = 48 *S*_0_ plants from KE being generated from the same seed batch as the DH-lines. Since *S*_0_ are heterozygous this corresponds to *n* = 96 haplotypes. Genotyping and quality control was performed in the same way as for the DH-lines, without heterozygosity filters. After imputation, an additional phasing step for the S0 using BEAGLE 4.1 (niterations=15) was performed. In both steps the DH-lines were used as a reference panel. Only markers overlapping with the DH dataset were included. This resulted in a second dataset containing *n* = 96 *S*_0_ and *n* = 501 DH haplotypes of KE and 487’462 markers.

Additionally, we used datasets from the 1000 Genomes Project phase 3 reference panel (1000 Genomes Project Consortium 2015) containing 5’008 haplotypes with a total of 88.3 million markers.

### Data Availability

The genetic data for maize, the associated R-package, the source code and a detailed documentation of the package is available at https://github.com/tpook92/HaploBlocker. Genetic data from the 1000 Genomes Project (1000 Genomes Project Consortium 2015) is available at ftp://ftp.1000genomes.ebi.ac.uk/vol1/ftp/release/20130502/. Supplemental files are available at FigShare.

## Results and Discussion

Here, we will focus on the results obtained for chromosome 1 (80’200 SNPs) of the landrace KE. All tests were also performed for all other chromosomes and the second landrace (PE) with similar results.

Using the previously described default settings of HaploBlocker, we identified 477 blocks which represent a coverage of 94.4% of the dataset and have an average length of 2’575 SNPs (median: 1’632 SNPs). For the whole genome, we identified 2’991 blocks representing 94.1% of the dataset with an average/median length of 2’685/1’301 SNPs. A graphical representation of the block structure for the first 20’000 markers of the set is given in Figure 5. Haplotypes were sorted according to their similarity at SNP 10’000. Since there is only limited linkage between markers further apart, the graphical representation gets increasingly fuzzy with increasing distance from the target SNP. For a comparison to a bifurcation plot (Sabeti et al. 2002; Gautier and Vitalis 2012) of that marker, we refer to the Supplementary Material (Figure S2).

**Figure 5.**
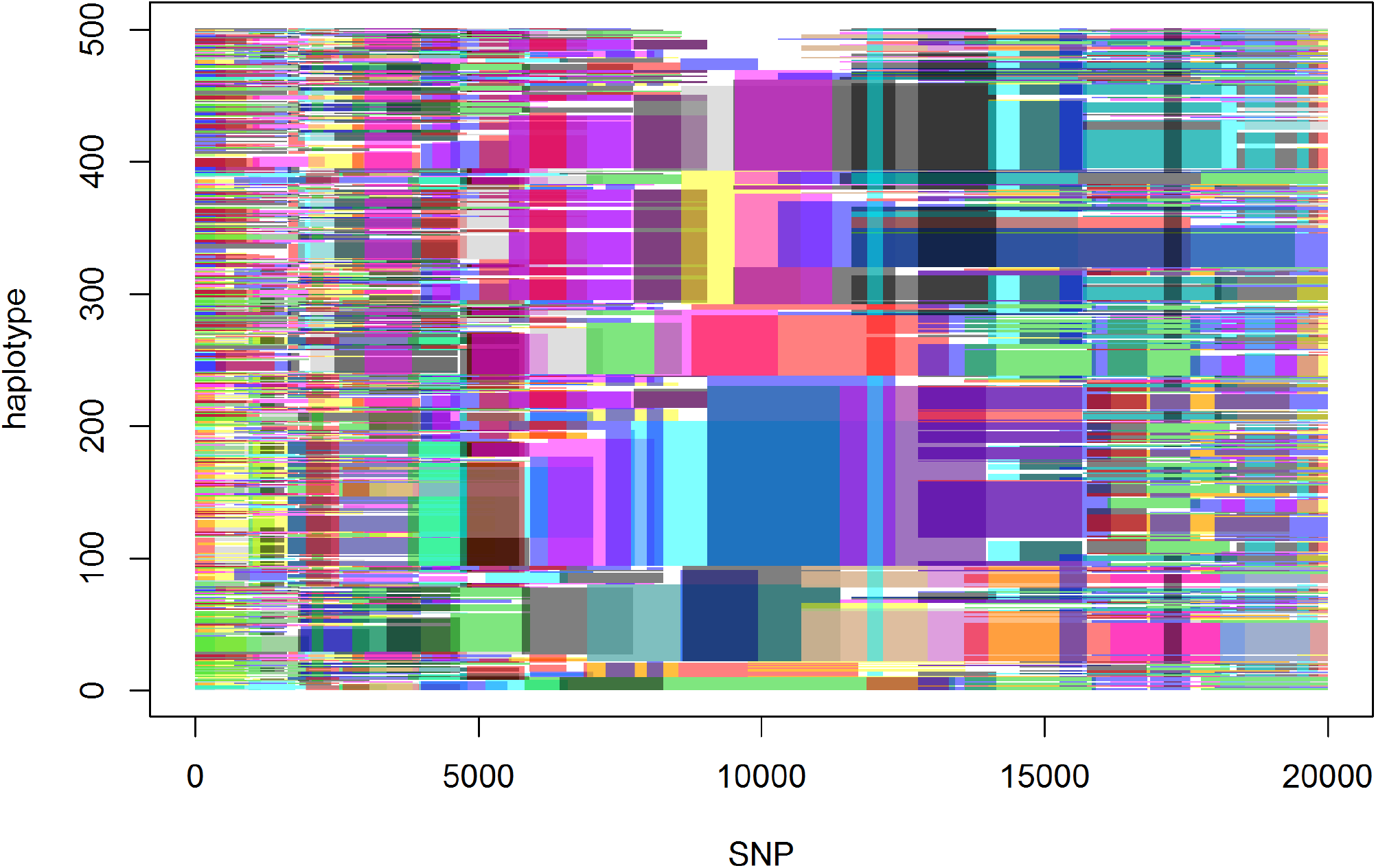
Graphical representation of the block structure for the first 20’000 SNPs of chromosome 1 in the KE DH-lines. Haplotypes are sorted for similarity in SNP 10’000. In that region block structures are most visible and transitions between blocks can be tracked easily. Further away from the centre the representation gets fuzzy.

When further investigating cells of the SNP-dataset that are not covered by any of the haplotype blocks, one can typically observe that in the associated segments the allelic sequence of the haplotype is a combination of multiple identified haplotype blocks, and by this indicating a recent recombination. Start and end points of blocks can be seen as candidates for positions of ancient (or at least non-recent) recombination, especially when multiple blocks start and end in the same region (e.g between markers 8’572 and 8’601 in Figure 5).

In the following, we will show and discuss the influence of certain parameter settings on the resulting haplotype library. Results will be evaluated according to the number of blocks, their length and the coverage of the haplotype library. Note that even though differences seem quite substantial, most haplotype libraries actually contain the same core set of haplotype blocks, which are the most relevant under basically any parameter setting. Parameter settings mostly influence which of the less relevant blocks are included. By this one can explicitly include a higher share of longer blocks, obtain a certain coverage or similar. For most routine applications, the use of the default settings with a target coverage should be sufficient.

### Effect of change in the MCMB

The MCMB affects both the number of blocks and the coverage of the dataset (Table 2). Higher MCMB leads to a stronger filtering of the haplotype blocks and thus to a haplotype library with lower coverage and decreased number of larger blocks. Overall, MCMB is the most important parameter to balance between conservation of information (coverage) and dimension reduction (number of blocks). It should be noted that the ideal parametrization of MCMB highly depends on data structure (e.g. marker density). Instead of using a set value for MCMB we recommend to fit the parameter automatically by setting a target coverage. For a graphical comparison of the structure of haplotype libraries with MCMB equal to 1’000, 5’000 and 20’000 we refer to the Supplementary Material (Figure S3).

**Table 2.** Influence of MCMB on the haplotype library for chromosome 1 in the KE DH-lines.

| MCMB | Number of Blocks | Average block length (# of SNPs) | Haplotypes per Block | Coverage |
| --- | --- | --- | --- | --- |
| 1 | 1'720 | 1'117 | 159.9 | 97.2% |
| 1'000 | 878 | 1'892 | 132.1 | 96.5% |
| 2'500 | 621 | 2'345 | 120.3 | 95.6% |
| 5'000 | 477 | 2'575 | 114.9 | 94.4% |
| 10'000 | 362 | 3'022 | 103.9 | 92.7% |
| 20'000 | 274 | 3'339 | 99.2 | 90.1% |
| 50'000 | 150 | 3'894 | 98.5 | 81.2% |

### Controlling length and number of haplotypes per block

The window size chosen in the cluster-building-step has a noteworthy influence on the window cluster. By using a smaller window size in the cluster-building-step, the resulting groups are bigger, leading to more and shorter (in terms of physical length) haplotype blocks in the block-identification-step (Table 3). As haplotype blocks are much larger than the window size in this case, the effects on the resulting haplotype library are only minor.

**Table 3.** Influence of the window size on the haplotype library for chromosome 1 in the KE DH-lines.

| Window size | Number of Blocks | Average block length (# of SNPs) | Haplotypes per Block | Coverage |
| --- | --- | --- | --- | --- |
| 5 | 488 | 2'489 | 121.8 | 93.5% |
| 10 | 482 | 2'544 | 113.7 | 93.6% |
| 20 | 477 | 2'575 | 114.9 | 94.4% |
| 50 | 474 | 2'615 | 101.4 | 95.0% |

In the block-filtering-step the weighting between segment length (*w_l_*) and number of haplotypes (*w_n_*) in each block influences the structure of the later obtained haplotype library (Table 4). As one would expect, a higher weighting for the length of a block leads to longer blocks that include fewer haplotypes. The effect of a lower relative weighting for the number of haplotypes in each block was found to have only a minor effect in our maize data. A possible reason for this is that even when using *w_l_* = *w_n_* the longest blocks previously identified were already selected in the haplotype library.

**Table 4.** Influence of the weighting of block length (*w_l_*) and number of haplotypes (*w_n_*) on the haplotype library for chromosome 1 in the KE DH-lines.

| $w_l$ | $w_n$ | Number of Blocks | Average block length (# of SNPs) | Haplotypes per Block | Coverage |
| --- | --- | --- | --- | --- | --- |
| 1 | 0 | 470 | 2'902 | 89.3 | 94.4% |
| 1 | 0.2 | 464 | 2'900 | 94.1 | 94.4% |
| 1 | 0.5 | 463 | 2'900 | 98.4 | 94.4% |
| 1 | 1 | 477 | 2'575 | 114.9 | 94.4% |
| 0.5 | 1 | 532 | 2'218 | 139.0 | 94.6% |
| 0.2 | 1 | 803 | 1'518 | 189.5 | 95.5% |
| 0 | 1 | 1313 | 934 | 208.2 | 96.1% |

When using the extended-block-identification method the average length of finally obtained haplotype blocks is massively increased in the obtained haplotype library (Table 5). Additionally, overlap between blocks is increased. Using this procedure will lead to the identification of the longest possible IBD segments, making it especially useful for applications like bEHH & IHH.

**Table 5.** Influence of using the extended-block-identification on the haplotype library in dependency of the parameter *t* of the extended-block-identification-step for chromosome 1 in the KE DH-lines.

| $t$ | Number of Blocks | Average block length (# of SNPs) | Haplotypes per Block | Coverage |
| --- | --- | --- | --- | --- |
| 1 | 477 | 2'575 | 114.9 | 94.4% |
| 0.95 | 603 | 5'659 | 89.8 | 94.6% |
| 0.9 | 788 | 9'371 | 70.2 | 95.2% |
| 0.8 | 916 | 11'716 | 60.9 | 95.5% |
| 0.6 | 970 | 12'430 | 58.5 | 95.7% |

Tests in this subsection were also performed when using a target coverage of 95%. For results on this we refer to the Supplemental Material (Table S2,S3,S4). Overall, results are similar.

### Haplotypes out of the sample

To assess how well HaploBlocker identifies haplotype block structures that also pertain to haplotype structures of other datasets, we split the maize data into a training and testing set and compared the share of both datasets represented by a haplotype library based on the training set alone. In all cases the coverage in the test set was below that of the training set, but with higher numbers of haplotypes in the training set the differences gets smaller. In case of 400 haplotypes in the training set and 101 haplotypes in the test set, the difference in coverage is as low as 2.7% (Figure 6) indicating that haplotype libraries derived in a sufficiently large dataset can be extended to individuals outside of the sample if they have similar genetic origin. Similar results were obtained when setting a target coverage (90%) for the test set.

**Figure 6.**
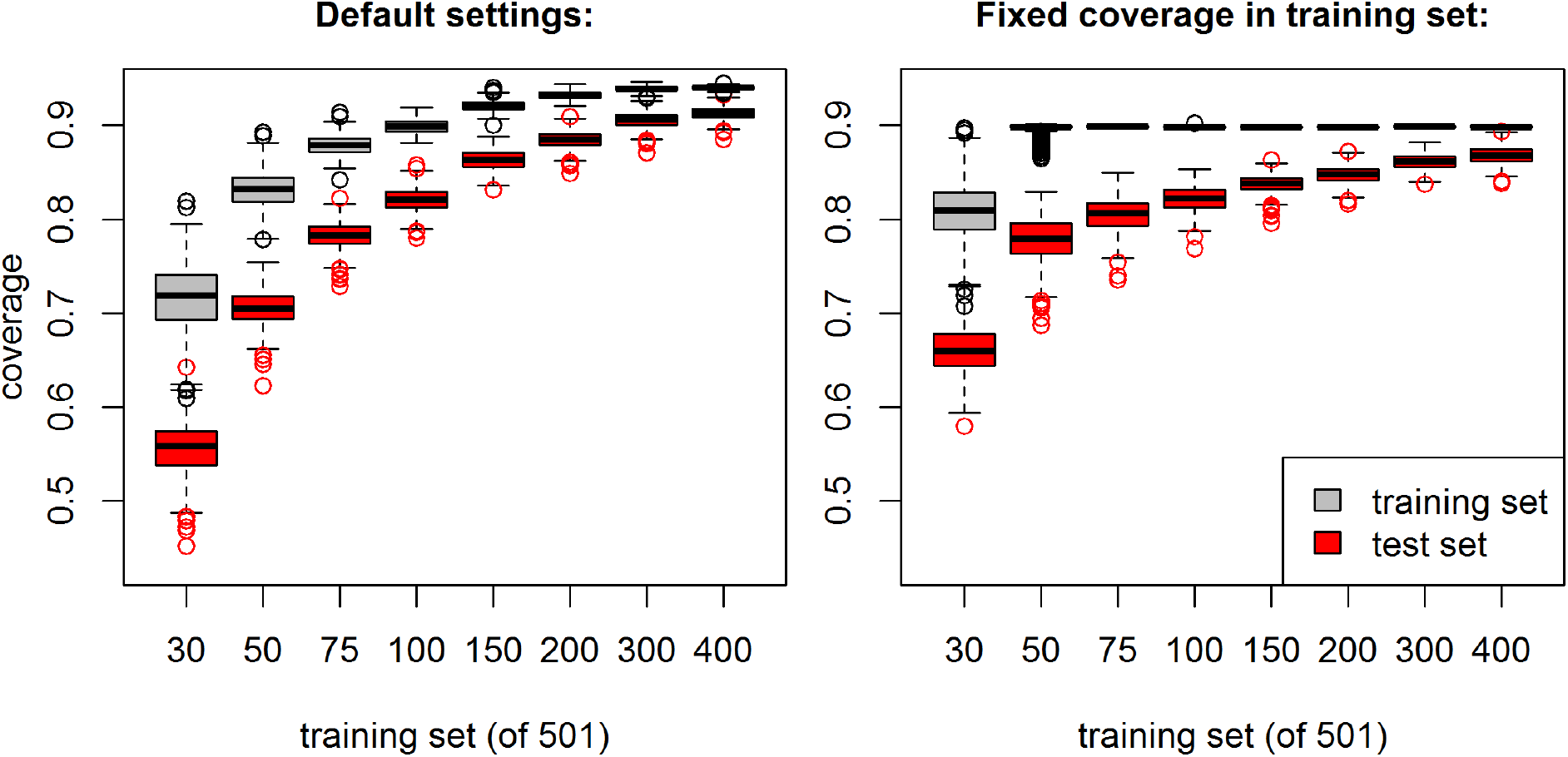
Proportion of the dataset represented by the haplotype library (coverage) of the training and test set in regard to size of the training set for chromosome 1 in the KE DH-lines.

### Information content

We investigated the information content between SNP- and block-dataset according to the method described above (de los Campos 2017), where *b_s_* was sampled from a standard Gaussian distribution. A REML approach was used for fitting the model. We found that, on average, 96.0% of the variance of the SNP-dataset can be explained by the default haplotype library (Table 6). As one would expect, the share of variance explained is increasing when increasing the number of blocks in the haplotype library. On the other hand, the share of the variance of the haplotype library that can be explained by the SNP-dataset is 95.2%. Even though the number of parameters in the block dataset (*Z*) is much smaller than in the full SNP set (*X*), the share of the variance explained by the respective other dataset is similar.

**Table 6.** Proportion of variance explained between the full SNP-dataset (*X*), a SNP-subset (*X_s_*) and the block dataset (*Z*). For comparability the number of parameters in *X_s_* and *Z* were chosen equally.

| Number of Blocks/SNPs | $X \sim Z$ | $Z \sim X$ | $X \sim X_s$ |
| --- | --- | --- | --- |
| 1'720 | 99.6% | 97.8% | 99.2% |
| 878 | 98.6% | 96.9% | 98.0% |
| 621 | 97.5% | 95.8% | 96.8% |
| 477 | 96.2% | 95.3% | 95.4% |
| 362 | 94.8% | 94.5% | 93.5% |
| 274 | 92.8% | 93.8% | 91.0% |
| 150 | 86.7% | 92.0% | 82.7% |

When using a subset of markers (*X_s_*) with the same number of SNPs as haplotype blocks in the haplotype library, the share of variation explained is slightly lower (95.1%) than for the block dataset. In contrary to the haplotype library, the variation of the SNP-subset is basically fully explained by the full SNP-dataset (≥ 99.99%). This should not be surprising since *X_s_* is a genuine subset of *X*. Even though a similar share in variation of the SNP-dataset is preserved, the block dataset should be preferred as it is able to incorporate effects that are not explained by linear effects of single markers.

With the following toy example, we illustrate what kind of effects can be grasped by a block dataset compared to a model that is only assigning effects to single markers, as is done in GBLUP (Meuwissen *et al.* 2001) using the traditional genomic relationship matrix (VanRaden 2008). Consider a dataset (Table 7) with three markers, five haplotypes and a genomic value of 1 for the allelic sequence 111. When assuming no environmental effects, phenotypes equal to genomic values and fitting an ordinary least squares model (OLS) on single markers, the resulting model estimates effects of 0.75, 0.5, and 0.5 for the three respective alleles with an intercept of −1. Overall, single marker effects can approximate but not fully explain an underlying epistatic genomic value (Table 7), whereas a block dataset allows for a natural model of effects caused by local interactions.

**Table 7.**
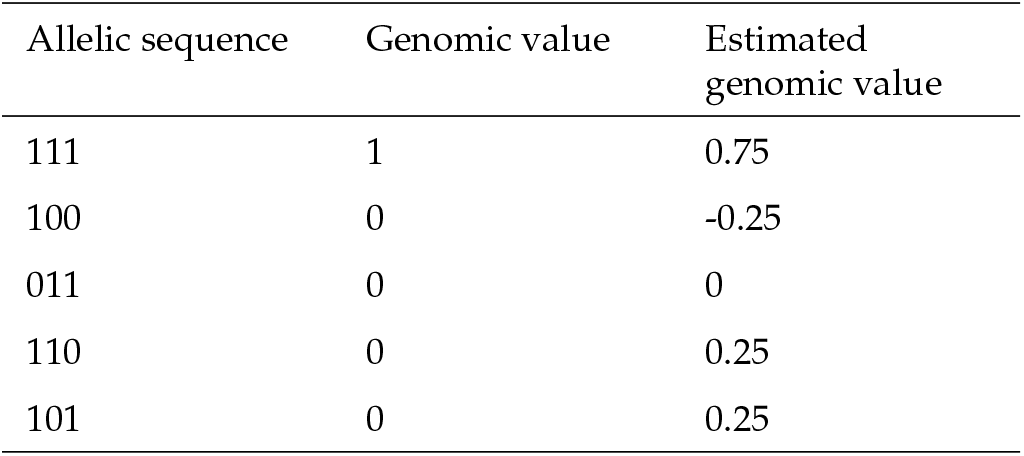
Estimated genomic values using an OLS model assuming additive effects of single markers.

| Allelic sequence | Genomic value | Estimated genomic value |
| --- | --- | --- |
| 111 | 1 | 0.75 |
| 100 | 0 | -0.25 |
| 011 | 0 | 0 |
| 110 | 0 | 0.25 |
| 101 | 0 | 0.25 |

### Overlapping segments in multiple landraces

When using HaploBlocker on the joint dataset of both landraces (KE & PE), the resulting haplotype library contains essentially the same haplotype blocks that were identified in the haplotype libraries derived for the two landraces individually. The reason for this is that segments shared between landraces are often short, leading to a small rating *r_b_* and thus removal in the block-filtering-step. To specifically identify those sequences that are present in both landraces, we added the constraint that each block had to be present in at least five haplotypes of both landraces. This results in the identification of 1’655 blocks which are present in both landraces. Those blocks are much shorter (avg. length: 207 SNPs) and represent only 62.7% of the genetic diversity of the dataset. This is not too surprising since the haplotypes of a single landrace are expected to be much more similar than haplotypes from different landraces. Explicitly, this is not an indicator for 62.7% of the chromosome of both landraces to be the same. Shared haplotype blocks can be found across the whole chromosome but only some haplotypes of the landraces have those shared segments.

### Comparison with the results of HaploView

Overall, the structure of the haplotype blocks generated with our approach is vastly different from blocks obtained with LD-based approaches such as HaploView (Barrett *et al.* 2005). When applying HaploView on default settings (Gabriel *et al.* 2002) to chromosome 1 of the maize data, 2’666 blocks are identified (average length: 27.8 SNPs, median: 20 SNPs) and 4’865 SNPs (6.1%) are not contained in any block. If one would use a similar coding to the blocks obtained in HaploBlocker and use a separate variable for each allelic sequence in a block, one would have to account for 12’550 different allelic sequences (excluding singletons). For the whole genome this would result in 16’904 blocks with 79’718 allelic sequences. When using a dataset with both landraces (or in general more diversity), LD-based blocks get even smaller (for chromosome 1: 4’367 blocks, 24’511 different allelic sequences, average length: 17.3 SNPs, median: 9 SNPs, 4’718 SNPs in no block). In comparison, the haplotype library identified in HaploBlocker with multiple landraces is, with minor exceptions, a combination of the two single landrace haplotype libraries (1’112 blocks, average length: 2’294 SNPs, median: 1’402 SNPs, coverage: 94.4%). Overall, the potential to detect long range associations between markers and to reduce the number of parameters in the dataset is much higher when using haplotype blocks generated by HaploBlocker.

Differences between the two methods become even more drastic when applying HaploView to the human datasets generated in the 1000 Genomes Project Phase 3 (1000 Genomes Project Consortium 2015). For chromosome 22 there were 49’504 blocks with an average length of 199 SNPs (median: 81 SNPs) that cover 92.9% of the dataset in HaploBlocker. In contrast, there were only 12’304 blocks (excluding singletons) identified in HaploView (average length: 8.1 SNPs, median: 4 SNPs) but only 99’130 of the 424’147 markers were assigned to a block (23.4%). In total, there were still 544’038 different allelic sequences in the identified blocks in HaploView. We noted that all alternative variants were coded as the same allele, as HaploView is only able to handle two alleles per marker, while HaploBlocker is able to handle up to 255 different alleles per marker. When allowing for more than two alleles per marker in HaploBlocker we obtain 49’500 blocks with an average length of 200 SNPs (median: 81 SNPs) that cover 93.0% of the dataset. It should be noted that HaploView was developed with different objectives in mind (Barrett *et al.* 2005).

### Influence of marker density

A common feature of conventional approaches to identify haplotype blocks is that with increasing marker density the physical size of blocks is strongly decreasing (Sun et al. 2004; Kim and Yoo 2016). To assess this, we executed HaploBlocker on datasets with different marker densities by only including every second/fifth/tenth/fortieth marker of the maize dataset in the model. Since the physical size of a window with a fixed number of markers is vastly different, we compared the structure of the obtained haplotype library using the adaptive mode in HaploBlocker (multiple window clusters with window sizes 5,10,20,50 and adaptive MCMB to obtain a target coverage of 95%) instead of default settings. As there are far fewer markers with possible variation, fewer blocks are needed to obtain the same coverage in the low-density datasets (Table 8). Since windows in the cluster-building-step span over a longer part of the genome, the considered groups contain fewer haplotypes leading to less frequent nodes in the window cluster. Since the haplotypes in a node are on average more related to each other, the identified blocks tend to be longer and include fewer haplotypes.

**Table 8.** Structure of the haplotype library under different marker densities using the adaptive mode in HaploBlocker with target coverage of 95% for chromosome 1 in the KE DH-lines.

| Density | Number of Blocks | Average block length<br>(# of SNPs on full array) | Haplotypes per Block | Used MCMB |
| --- | --- | --- | --- | --- |
| Every SNP | 534 | 2'317 | 116.4 | 2'813 |
| Every second SNP | 523 | 2'281 | 112.7 | 1'563 |
| Every fifth SNP | 450 | 2'557 | 96.9 | 945 |
| Every tenth SNP | 401 | 2'811 | 90.6 | 758 |
| Every fortieth SNP | 319 | 3'637 | 79.9 | 294 |

In a second step, we manually adapted the window size (50/25/10/5/5) and the MCMB (5000/2500/1000/500/125) according to the marker density of the dataset. When manually adapting the parameters, the number of blocks in the haplotype library is largely independent of the marker density (Table 9). The length of the blocks is decreasing, whereas the number of haplotypes per block is increasing with decreasing marker density. A possible reason for this is that haplotypes in the same node of the window cluster are less similar in the region than when using bigger window sizes. This will lead to shorter haplotype blocks which are carried by more but less related haplotypes. In case of the dataset in which we used every fortieth marker, we additionally considered a value of 250 for the MCMB since the resulting coverage was a lot higher, indicating that less overall variation is present in the dataset. This also results in fewer overall blocks needed to obtain similar coverage.

**Table 9.**
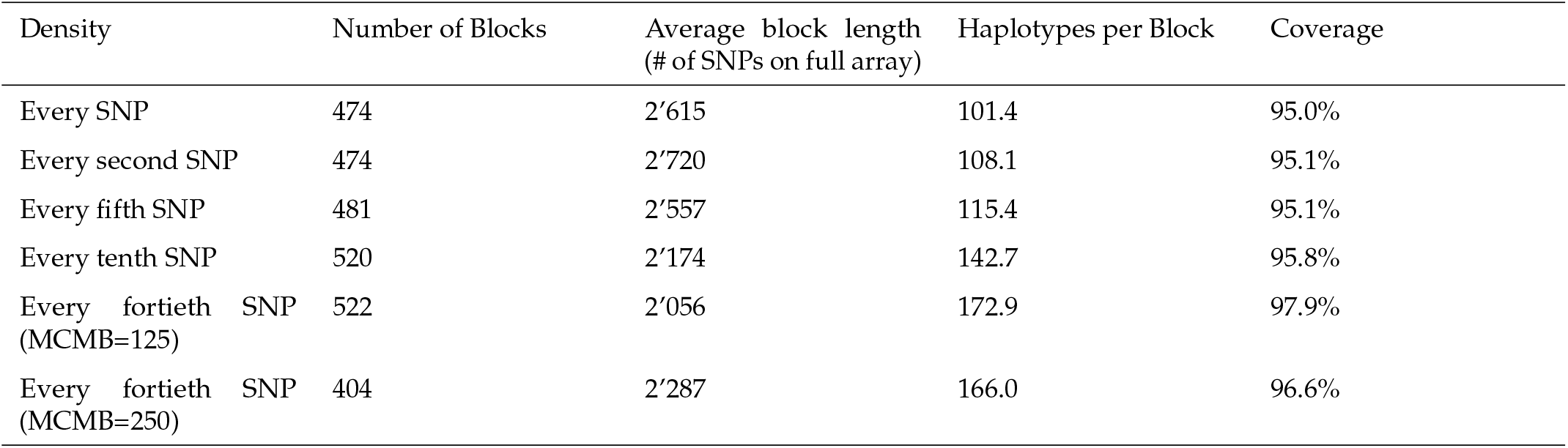
Structure of the haplotype library under different marker densities when adjusting parameters according to data structure for chromosome 1 in the KE DH-lines.

Haplotype libraries for all considered marker densities were similar, indicating that for our landrace population a much lower marker density would have been sufficient to derive haplotype blocks via HaploBlocker. In case the physical size of haplotype blocks is smaller, a higher marker density is needed.

### Recovering founder haplotypes

HaploBlocker was applied to the final generation of the dataset simulated in analogy to the breeding scheme for the MAGIC population given in (Zheng *et al.* 2015). On average, we obtained 827 haplotype blocks with a length of 1’458 markers covering 82.6% of the dataset. 96.6% of the allelic sequences of haplotype blocks are at least 99.5% the same as an allelic sequence of a founder haplotype of that segment. Overall 86.1% of all cells in the SNP-dataset of the founders are recovered by the resulting haplotype library. When using a target coverage of 95%, the share of the allelic sequences of the blocks that are the same as a founder haplotype are quite similar (96.9%) but 92.4% of all cells of the SNP-dataset of the founders can be recovered. Note that identified haplotype blocks, on default, have a minimum size of 5 haplotypes, leading to the loss of rarely inherited haplotypes. It should be noted that our approach is not constructed to detect the exact boundaries of IBD segments between founders and single offspring but instead is detecting commonly presented allelic sequences. In a population with limited founders (e.g. caused by a genetic bottleneck), those common allelic sequences most likely stem from the founders of the population. For a plot comparing the true and estimated genetic origin of the final generation we refer to Figure 7. Here, estimation means that in case a haplotype block completely stems from a single founder that particular founder is used as the origin. In practice non-overlapping blocks can of course not be assigned to the same founder. Main benefit of our method is that in contrast to commonly used methods only phased genotype data is needed to recover founder haplotypes. When interest is in the exact boundaries of IBD segments for single haplotypes and founders (with known pedigree), we recommend the usage of methods like RABBIT (Zheng *et al.* 2015).

**Figure 7.**
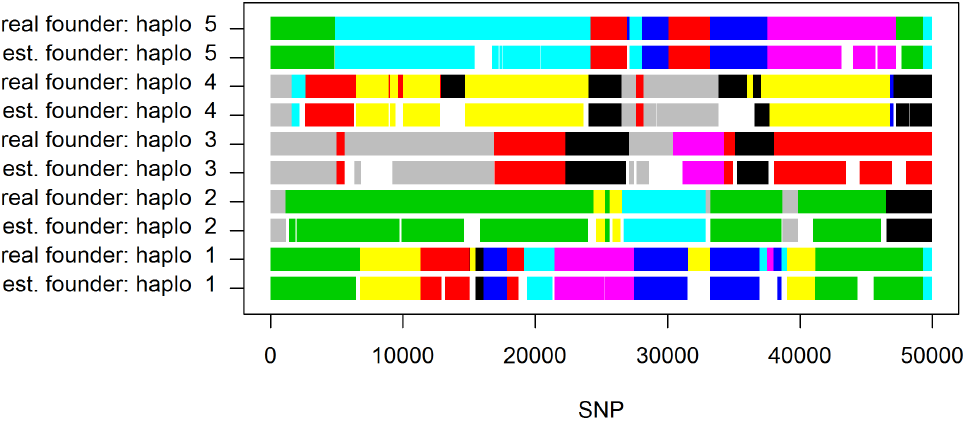
Estimated and true founders for the first five haplotypes of the last generation of a MAGIC population simulated according to breeding scheme given in (Zheng *et al.* 2015) using a target coverage of 95% in the generation of the haplotype library.

### Block-based selection signatures

When deriving EHH and bEHH scores, we can observe that curves are quite similar for DH-lines (Figure 8). Most apparent difference is a much higher EHH score in the directly surrounding region of the marker. Those segments are typically much smaller than the segments considered jointly in the bEHH approach. Note that the same allelic sequence in such a small region can not only occur based on IBD but also by chance. On the contrary, scores between distant markers for the *S*_0_ plants are much lower when using EHH (Figure 8). This is mainly caused by the incorporated robustness of bEHH, since the S0 dataset tends to contain a higher share of minor deviations between haplotypes.

**Figure 8.**
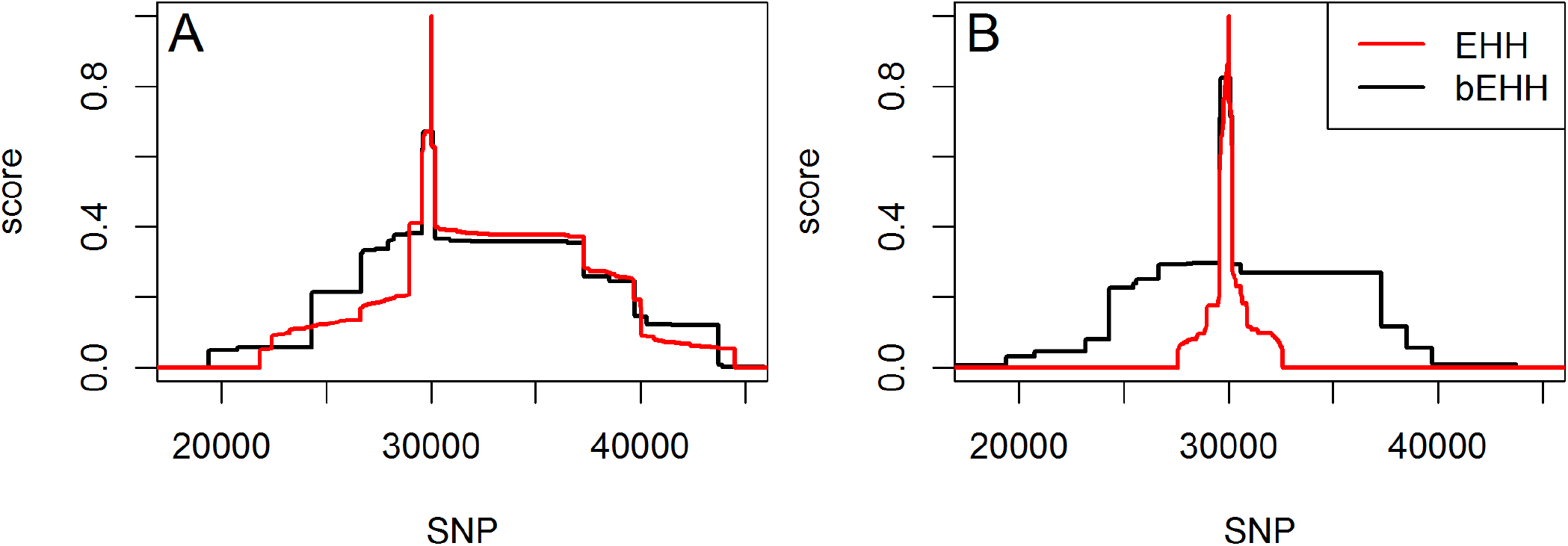
Comparision of EHH and bEHH scores for DH-lines (A) and *S*_0_ (B) for marker 30’000 of chromosome 1 in the KE DH-lines.

When using EHH (Sabeti *et al.* 2002) to derive IHH (Voight *et al.* 2006), the selection pressure on DH-lines is estimated to be much higher, whereas scores are quite similar between the two groups when using bEHH (Figure 9). IHH scores based on bEHH are in concordance with previous research, as we would expect little to no loss of diversity or selection in the process of generating DH-lines (Melchinger *et al.* 2017). Results in Melchinger *et al.* (2017) were derived by the use of *F_st_* (Holsinger and Weir 2009) and analysis of molecular diversity in single markers. As presented at a recent conference (Mayer *et al.* 2018), similar studies with matching results were also performed for KE and PE.

**Figure 9.**
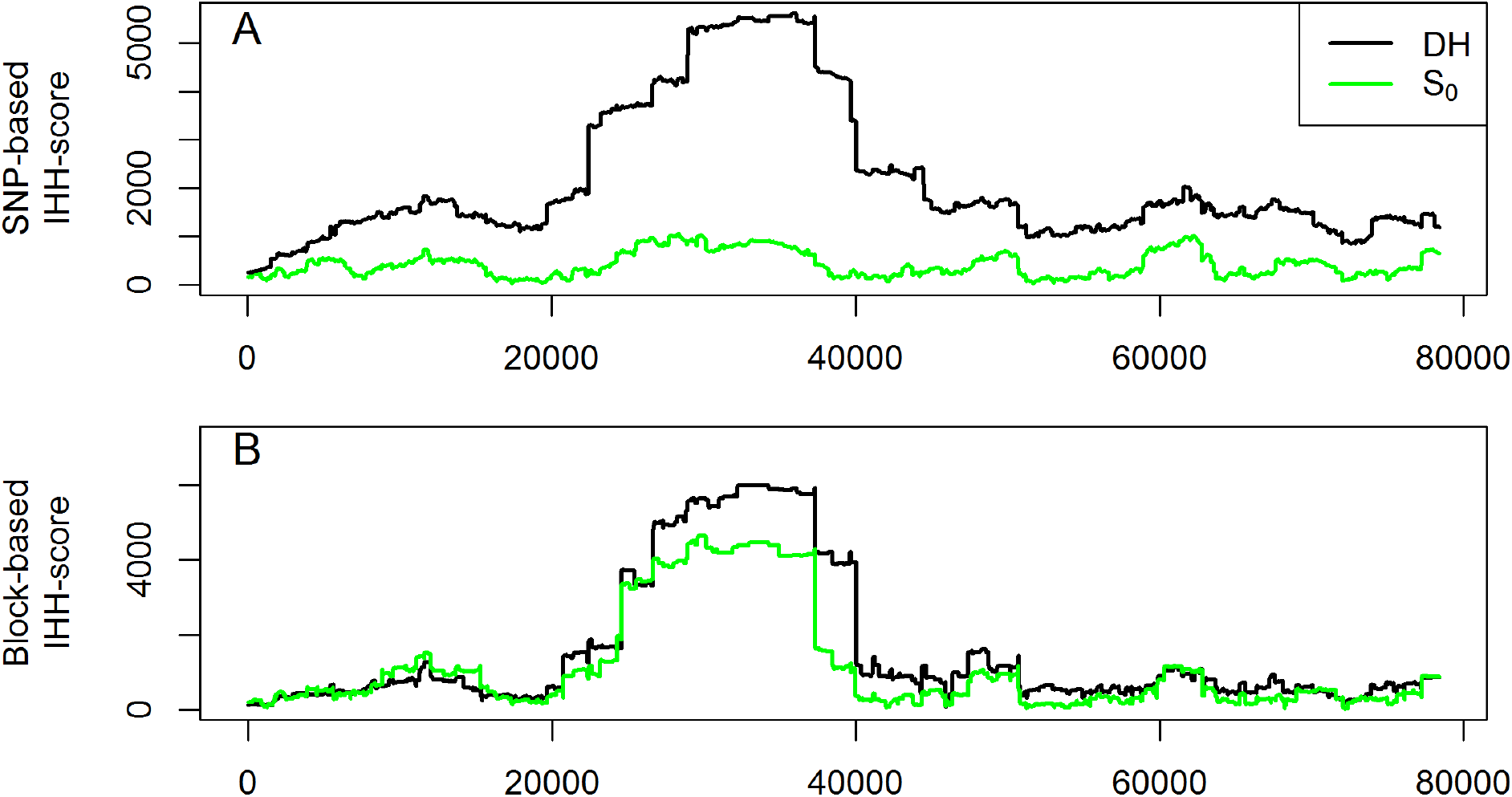
IHH scores based on SNPs (A) and haplotype blocks (B) for DH-lines and S0 for chromosome 1 in the KE.

### Computing time

Overall computing times were not an issue for the considered datasets when using the associated R-package HaploBlocker (R Core Team 2017; Pook and Schlather 2018) with the full dataset (501 haplotypes, 80’200 SNPs) needing 55 seconds on default, 75 seconds with a target coverage and 13.3 minutes in the adaptive mode. Computations were performed on a single core of a server cluster with Broadwell Intel E5-2650 (2X12 core 2.2 GHz) processors. Most crucial parts in terms of computing time are written in C.

For our datasets, computing time scaled approximately linear in both the number of haplotypes and the physical size of the genome analyzed (Figure 10). Especially for the number of haplotypes it is difficult to generalize because the number of nodes in the window cluster is mainly causal for the increase in computing time. The marker density only had a minor effect. Even a panel containing just every tenth marker, on average, needed 99.3% of the computing time of the full dataset.

**Figure 10.**
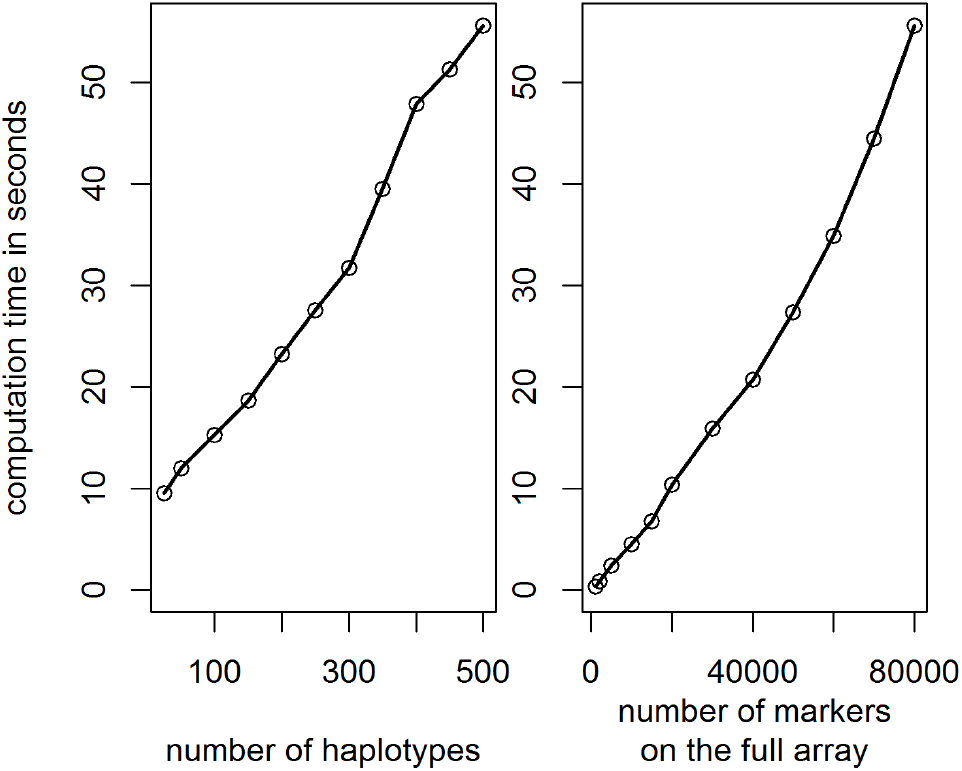
Comparison of computing times for datasets of various sizes for chromosome 1 in the KE DH-lines.

### Conclusions and Outlook

HaploBlocker provides a natural technique to model local epistasis and thereby solves some of the general problems of markers being correlated but not causal individually (He et *al.* 2017; Akdemir et *al.* 2017). This can be seen as one of the factors contributing to the “missing heritability” phenomenon in genetic datasets (Manolio et *al.* 2009).

Even though results were mainly presented for a maize dataset, methods are not species-dependent and were also applied on livestock and human data. However, the opportunities for identifying long shared segments will be higher in SNP-datasets from populations subjected to a recent history of intensive selection. When using data containing individuals with heterozygous loci a high phasing accuracy is needed since HaploBlocker is not able to handle uncertainty in haplotype phase assignment. Especially for human data this can be a substantial problem in the application and thereby requiring triplet data or high quality phase like in the 1000 Genomes Project (1000 Genomes Project Consortium 2015).

It should be noted that by using blocks, an assignment of effects to physical positions (like in a typical GWAS study) is not obtained. A subsequent analysis is needed to identify which segment of the significantly trait-associated haplotype block is causal for a trait and/or which parts of that block differ from the other blocks in that region.

A future topic of research is the explicit inclusion of larger structural variation like duplications, insertions or deletions as is done in methods to generate a pangenome (Eggertsson *et al.* 2017). Since blocks in HaploBlocker are of large physical size most structural variation should still be modelled implicitly and an application to sequence data is perfectly possible. HaploBlocker provides an innovative and flexible approach to screen a dataset for block structure. The representation and condensation of a SNP-dataset as a block dataset is enabling new methods for further genomic analyses. For some applications, already existing techniques for a SNP-dataset can directly be applied by using a block dataset instead (e.g. genomic prediction). For other applications, like the detection of selection signatures via EHH/IHH, modifications of the original methodology are needed. Features of HaploBlocker can even enhance existing methods and lead to improvements like an increased robustness of the methods against minor variation or a massively reduced computing time. Additionally, problems regarding typical *p* ≫ *n*–settings in genetic datasets (Fan *et al.* 2014) can be heavily reduced, allowing for the usage of more complex statistical models that include epistasis or even apply deep learning methods with a reduced risk of over-fitting.

## Acknowledgments

The authors thank the German Federal Ministry of Education and Research (BMBF) for the funding of our project (MAZE - “Accessing the genomic and functional diversity of maize to improve quantitative traits”; Funding ID: 031B0195). Gustavo de los Campos was supported by a Mercator-fellowship of the German Research Foundation (DFG) within the Research Training Group 1644, “Scaling problems in statistics” (grant no. 152112243). We further thank Nicholas Schreck for useful comments on the manuscript and his help regarding unbiased heritability estimation.

---

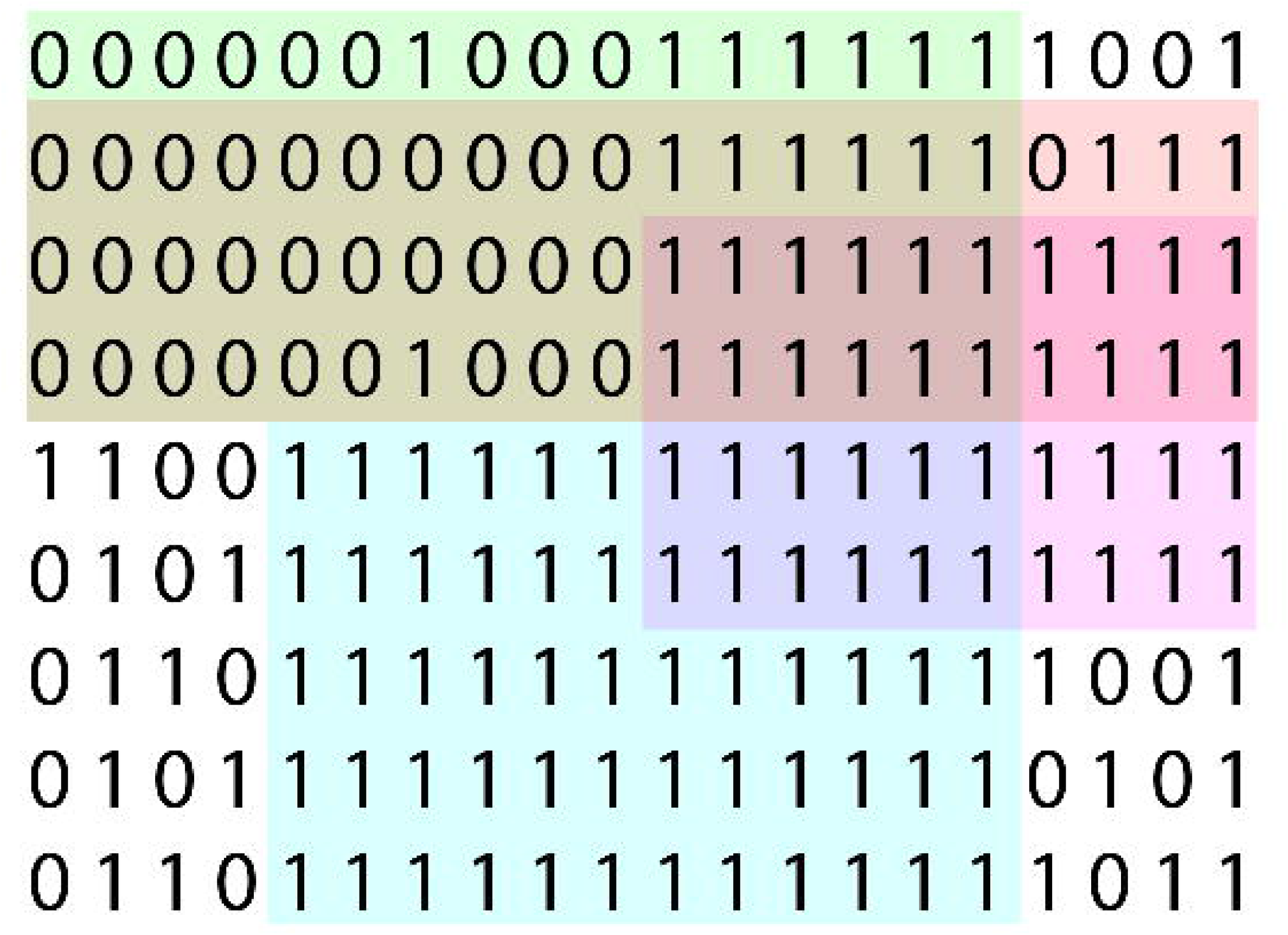

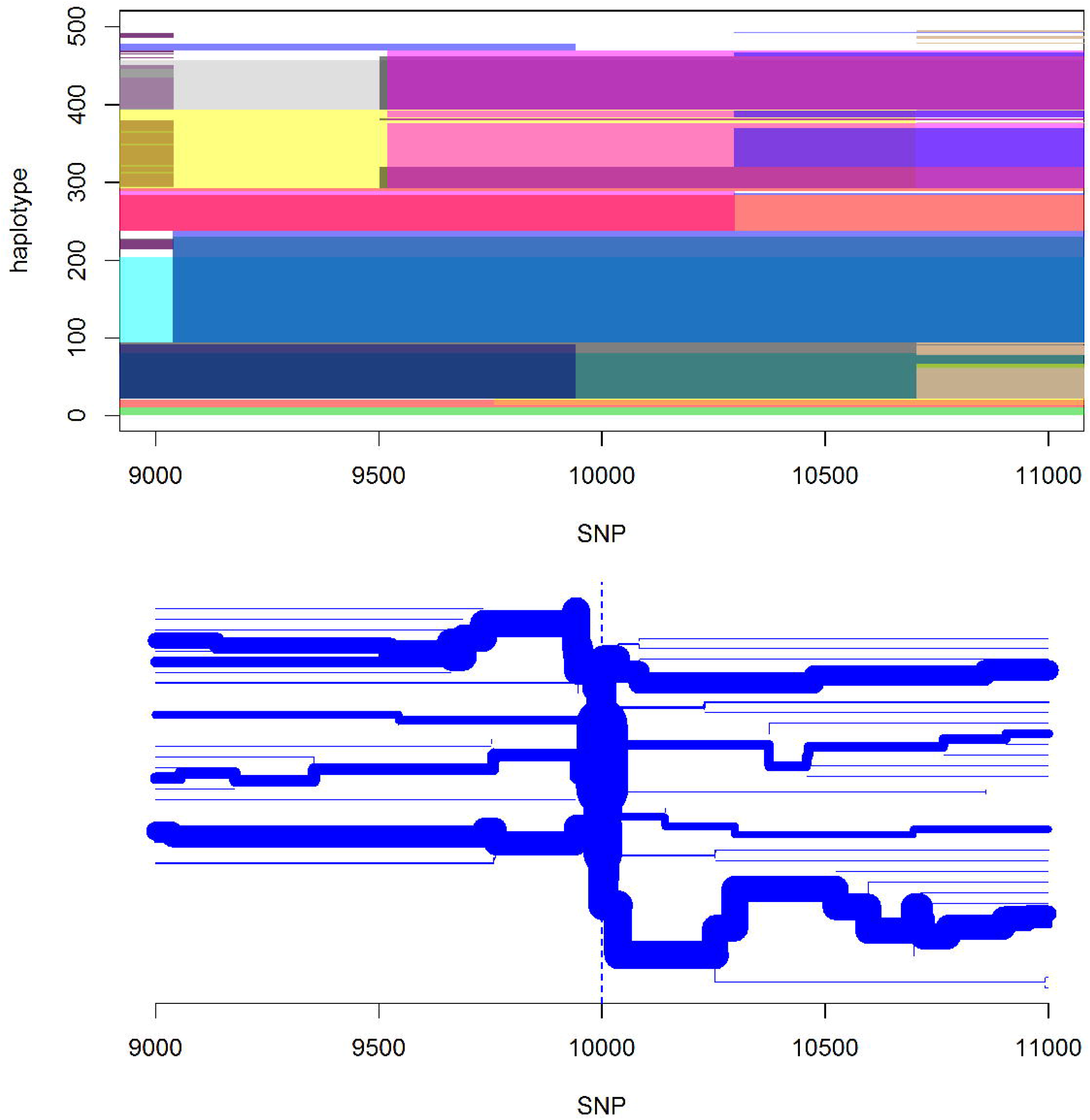

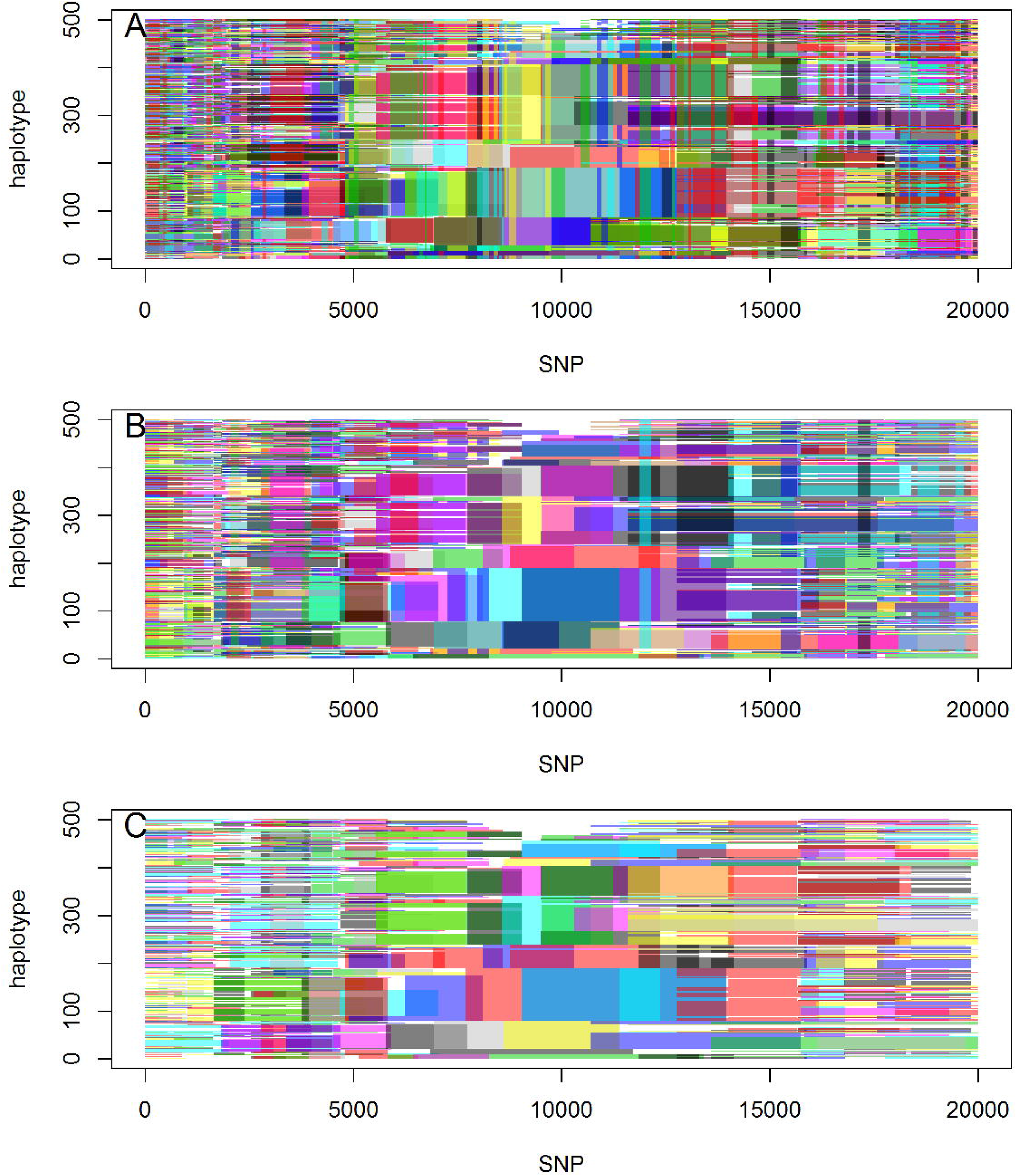

