## Supplemental File 1 for "HaploBlocker: Creation of subgroup specific haplotype blocks and libraries"

**Target-coverage (optional)**

In the following, we will denote the share of the dataset that is represented by the haplotype library as the coverage of the dataset. To control the coverage, we propose an adaptive fitting of the MCMB. Especially for different marker densities the choice of the MCMB is relevant to control the minimum size of each block and thus the resulting obtained coverage. The MCMB is fitted by iteratively increasing/decreasing the MCMB when the coverage is too high/low. We double/halve the value of the MCMB from step to step until both a haplotype library with a higher and lower coverage than the target exists. Afterwards the mean of the MCMB values of the two libraries with coverage closes (one above/below) to the target are used next. This procedure is repeated until the MCMB is 1 or the target coverage is reached.
