## Supplemental File 2 for "HaploBlocker: Creation of subgroup specific haplotype blocks and libraries"

**Graphical representation of haplotype blocks**

We suggest a graphical representation of haplotype blocks to show transition rates between blocks in analogy to bifurcation plots (Sabeti et al., 2002; Gautier and Vitalis, 2012). To this end, we first sort the blocks of the haplotype library according to the physical position of the first SNP of the block. In case of identical starting points the shorter block is considered first. Our aim in sorting the haplotypes is to cluster haplotypes according to their similarity around a specific physical position (default: SNP in the middle of the dataset). The sorting process itself is executed in two alternating steps:

**Step 1: Adding new haplotypes**

In the first iteration of this step we select all haplotypes in the most common block that includes the marker we want to align against. In later iterations, we add the haplotypes of the block with the biggest overlap of haplotypes with the previously considered block. In case no block has any overlapping haplotypes, the block with the most haplotypes not considered so far is used next.

**Step 2: Sorting new haplotypes**

The newly added haplotypes are ordered according to their presence in physically close blocks. We do this by iteratively comparing the haplotypes of other blocks, starting with the directly adjacent ones. Whenever only some of the currently considered haplotypes are in an adjacent block, we split haplotypes into two groups and proceed with both groups separately. This procedure is stopped when every group has either exactly one haplotype left or the end of the haplotype library has been reached.
