## Supplemental File 3 for "HaploBlocker: Creation of subgroup specific haplotype blocks and libraries"

**Toy example bEHH**

As a toy example consider a dataset with 9 haplotypes and 4 haplotype blocks (Supplementary Material Figure S1): green (markers: 1-16), blue (5-16), red (1-20) and purple (11-20). For simplicity we are using marker numbers as physical positions in this example. Resulting block boundaries to consider are 1,5,11,17,21.

EHH between markers 1 and 4 is higher than bEHH for the segment:

$$bEHH\left( \left[ 1,4 \right],\left[ 1,4 \right] \right)= \frac{\binom{4}{2}}{\binom{9}{2}}= \frac{1}{6}$$

$$EHH\left( 1,4 \right)= \frac{\binom{4}{2}+\binom{2}{2}+\binom{2}{2}}{\binom{9}{2}}= \frac{2}{9}$$

The higher score is caused by the allelic sequences 0101 and 0110 that both occur twice. Those haplotypes are not part of a haplotype block and thus are ignored in bEHH. For both scores the allelic sequence 0000 and the associated haplotype are present four times.

When considering EHH between markers 5 and 16 or segments [5,10] and [11,16] following scores are obtained:

$$bEHH\left( \left[ 5,10 \right],\left[ 11,16 \right] \right)= \frac{\binom{4}{2}+\binom{5}{2}}{\binom{9}{2}}= \frac{4}{9}$$

$$EHH\left( 5,16 \right)= \frac{\binom{2}{2}+\binom{2}{2}+\binom{5}{2}}{\binom{9}{2}}= \frac{1}{3}$$

Allelic sequences 00100111111 and 00000111111 are considered jointly in bEHH, as they are part of the same haplotype block. For EHH, the two allelic sequences are considered separately. Note that this is toy example and in reality blocks are much longer. Haplotypes that are not contained in any block can be considered separated. As this massively increases computing time, this is not done by default.
